# Baseline T cell immune phenotypes predict virologic and disease control upon SARS-CoV infection

**DOI:** 10.1101/2020.09.21.306837

**Authors:** Jessica B. Graham, Jessica L. Swarts, Sarah R. Leist, Alexandra Schäfer, Vineet D. Menachery, Lisa E. Gralinski, Sophia Jeng, Darla R. Miller, Michael A. Mooney, Shannon K. McWeeney, Martin T. Ferris, Fernando Pardo-Manuel de Villena, Mark T. Heise, Ralph S. Baric, Jennifer M. Lund

**Affiliations:** Vaccine and Infectious Disease Division, Fred Hutchinson Cancer Research Center, Seattle, WA; Department of Epidemiology, University of North Carolina at Chapel Hill, Chapel Hill, NC; Department of Microbiology and Immunology, University of Texas Medical Center, Galveston, TX; OHSU Knight Cancer Institute, Oregon Health & Science University, Portland, OR; Oregon Clinical and Translational Research Institute, Oregon Health & Science University, Portland, OR; Department of Genetics, University of North Carolina at Chapel Hill, Chapel Hill, NC; Lineberger Comprehensive Cancer Center, University of North Carolina at Chapel Hill, Chapel Hill, NC; Division of Bioinformatics and Computational Biology, Department of Medical Informatics and Clinical Epidemiology, Oregon Health & Science University, Portland, OR; Department of Global Health, University of Washington, Seattle, WA

**Keywords:** SARS-CoV, immune correlates of disease, Collaborative Cross

## Abstract

The COVID-19 pandemic has revealed that infection with SARS-CoV-2 can result in a wide range of clinical outcomes in humans, from asymptomatic or mild disease to severe disease that can require mechanical ventilation. An incomplete understanding of immune correlates of protection represents a major barrier to the design of vaccines and therapeutic approaches to prevent infection or limit disease. This deficit is largely due to the lack of prospectively collected, pre-infection samples from indiviuals that go on to become infected with SARS-CoV-2. Here, we utilized data from a screen of genetically diverse mice from the Collaborative Cross (CC) infected with SARS-CoV to determine whether circulating baseline T cell signatures are associated with a lack of viral control and severe disease upon infection. SARS-CoV infection of CC mice results in a variety of viral load trajectories and disease outcomes. Further, early control of virus in the lung correlates with an increased abundance of activated CD4 and CD8 T cells and regulatory T cells prior to infections across strains. A basal propensity of T cells to express IFNg and IL17 over TNFa also correlated with early viral control. Overall, a dysregulated, pro-inflammatory signature of circulating T cells at baseline was associated with severe disease upon infection. While future studies of human samples prior to infection with SARS-CoV-2 are required, our studies in mice with SARS-CoV serve as proof of concept that circulating T cell signatures at baseline can predict clinical and virologic outcomes upon SARS-CoV infection. Identification of basal immune predictors in humans could allow for identification of individuals at highest risk of severe clinical and virologic outcomes upon infection, who may thus most benefit from available clinical interventions to restrict infection and disease.

**Summary:** We used a screen of genetically diverse mice from the Collaborative Cross infected with mouse-adapted SARS-CoV in combination with comprehensive pre-infection immunophenotyping to identify baseline circulating immune correlates of severe virologic and clinical outcomes upon SARS-CoV infection.

## Introduction

The SARS-CoV-2 pandemic has led to a massive number of infections worldwide, with an unprecedented combined toll in terms of mortality, long-term health conditions, and economic turmoil (Dong et al., 2020). While large-scale efforts to develop protective vaccines are underway, the human immune response to natural infection and identification of immune correlates of disease outcome and protection is still in process. These efforts are likely to help guide such vaccine efforts, as an understanding of the natural immune correlates of protection from disease could assist in the rational design of prophylactic or therapeutic vaccines against SARS-CoV-2, as well as potential immunotherapeutic strategies. Multiple studies have demonstrated that following infection with SARS-CoV-2, individuals can present with mild or asymptomatic disease, though a subset of patients experience severe disease that often requires hospitalization and ventilation. Thus, some of the first studies of the human immune response to SARS-CoV-2 infection have examined changes in immune cell populations in peripheral blood from patients with severe disease as compared to healthy controls. Such studies of patients with severe COVID-19 have identified the existence of SARS-CoV-2-specific CD4 and CD8 T cells (Grifoni et al., 2020; Mateus et al., 2020; Weiskopf et al., 2020), as well as an interferon-stimulated gene signature (Wilk et al., 2020), and various changes in immune cell dynamics (Lucas et al., 2020; Mathew et al., 2020; Wilk et al., 2020). Notably, most studies have reported dysregulated and/or inflammatory responses in patients with severe COVID-19, including decreases in regulatory T cells (Qin et al., 2020), increased neutrophil counts (Lucas et al., 2020; Qin et al., 2020; Wilk et al., 2020) and increases in pro-inflammatory cytokines such as IL-6 and TNF (Blanco-Melo et al., 2020; Lucas et al., 2020; Qin et al., 2020), thereby suggesting that a dysregulated state of inflammation is associated with severe COVID-19. However, what is thus far lacking is a study of prospectively collected, pre-infection samples that would serve to identify if there are immune correlates of protection from infection and/or from severe disease upon infection with SARS-CoV-2. Because most studies have been conducted after individuals had been infected with SARS-CoV-2, it is unclear if the identified immune signatures are predictive of severe disease or a manifestation of severe disease.

Previous studies of immunity to other coronaviruses have also contributed to our understanding of what to expect from SARS-CoV-2 in terms of immunity (Sariol and Perlman, 2020). Specifically, studies of samples from survivors of MERS-CoV infection have determined that the development of CD4+ and CD8+ T cell responses occurs in humans (Zhao et al., 2017), and studies of SARS-CoV and MERS-CoV infection in mice have demonstrated that protection is mediated by airway memory CD4+ T cells (Zhao et al., 2016). Given this published evidence from human infection with SARS-CoV-2 plus these studies of other CoVs demonstrating that T cells are likely to be involved in immunity to CoV infections, we reasoned that it is possible that T cells could play a role in the initial stages of infection, and thus a pre-infection assessment of the T cell phenotype could reveal novel predictors of severe virologic and clinical outcomes upon infection. Further, given that a dysregulated, pro-inflammatory state is associated with severe COVID-19, we hypothesized that such a signature prior to infection might be predictive of disease outcome upon infection.

Immune correlates in humans are normally difficult to identify as they require a prospective, longitudinal study of immune responses in infected individuals pre- and post-infection. Animal models, on the other hand, have many advantages, such as the ease of study of immunity at pre- and post-infection timepoints, as well as experimental control over most variables including timing of infection, infection dose, host genetics, diet, and infection route. Therefore, we have used the Collaborative Cross (CC), a population of genetically diverse, recombinant inbred mouse strains, to investigate whether pre-infectious immune predictors were related to SARS-CoV disease. CC strains are derived from eight founder mouse strains that include five classical inbred strains and three wild-derived strains using a funnel breeding strategy followed by inbreeding (Churchill et al., 2004; Collaborative Cross, 2012; Keane et al., 2011; Roberts et al., 2007b). It is well-documented that the CC can be used to model the diversity in human immune responses and disease outcomes that are not present in standard inbred mouse models (Brinkmeyer-Langford et al., 2017; Elbahesh and Schughart, 2016; Ferris et al., 2013; Graham et al., 2018; Graham et al., 2016; Graham et al., 2015; Gralinski et al., 2015; Kollmus et al., 2018; Leist and Baric, 2018; Rasmussen et al., 2014). We have previously shown that the CC is a superior model for the vast diversity in T cell phenotypes present in the human population (Graham et al., 2017b), and also used a screen of F1 mice derived from CC crosses (CC recombinant intercross, CC-RIX) infected with three different RNA viruses (H1N1 influenza A virus, SARS-CoV, and West Nile virus) to reveal novel baseline immune correlates that are associated with protection from death upon infection from all of these three viruses (Graham et al., 2020). Here, we focus our analysis on specific circulating, pre-infection immune phenotypes that associate with different virologic and clinical outcomes upon SARS-CoV infection, including uncontrolled virus replication in the lung, weight loss, and death. We find evidence to support the notion that a circulating dysregulated and inflammatory immunophenotype prior to infection is associated with severe virologic and clinical disease outcomes upon infection with SARS-CoV. While further testing in animal modes and humans is required, our data are consistent with the notion that a test of circulating immune signatures could be used to predict infection outcomes and thereby identify patients at highest risk of high rates of shedding and disease upon infection that would most benefit from targeted therapeutic interventions.

## Methods

### Mice

CC mice were obtained from the Systems Genetics Core Facility at the University of North Carolina-Chapel Hill (UNC) (Welsh et al., 2012). As reported previously (Graham et al., 2020), between 2012 and 2017, F1 hybrid mice derived from intercrossing CC strains (CC-RIX) were generated for this research study at UNC in an SPF facility based on the following principles: (1) Each CC strain used in an F1 cross had to have been certified distributable (Welsh et al., 2012); (2) The UNC Systems Genetics Core Facility was able to provide sufficient breeding animals for our program to generate N=100 CC-RIX animals in a target three month window; (3) Each CC-RIX had to have one parent with an *H2B^b^* haplotype (from either the C57BL/6J or 129S1/SvImJ founder strains), and one parent with a haplotype from the other six CC founder strains; (4) Each CC had to be used at least once (preferably twice) as a dam, and once (preferably twice) as a sire in the relevant CC-RIX; (5) Lastly, we included two CC-RIX multiple times across the five years of this program to specifically assess and control for batch and seasonal effects. The use of CC-RIX allowed us to explore more lines than the more limited number of available RI strains, and additionally, CC-RIX lines were bred to ensure that lines were heterozygous at the H-2b locus, having one copy of the H-2b haplotype and one copy of the other various haplotypes. This design was selected such that we could examine antigen-specific T-cell responses for our parallel studies of immunogenetics of virus infection, while concurrently maintaining genetic variation throughout the rest of the genome.

Six to eight week old F1 hybrid (RIX) male mice were transferred from UNC to the University of Washington and housed directly in a BSL-2+ laboratory within an SPF barrier facility. Concurrelty, F1 hybrid female mice were transferred internally to UNC to a BSL-3 facility for SARS-CoV infection. Male 8-10 week old mice were used for all baseline immune experiments, with 3-6 mice per experimental group. All animal experiments were approved by the UW or UNC IACUC. The Office of Laboratory Animal Welfare of NIH approved UNC (#A3410-01) and the UW (#A3464-01), and this study was carried out in strict compliance with the PHS Policy on Humane Care and Use of Laboratory Animals.

### Virus and Infection

Mouse adapted SARS-CoV MA15 (Roberts et al., 2007a) was propagated and titered on Vero cells as previously described (Gralinski et al., 2015; Gralinski et al., 2018). For virus quantification from infected mice, plaque assays were performed on lung (post-caval lobe) tissue homogenates as previously described (Gralinski et al., 2017). Mice were intranasally infected with 5×10^3^ PFU of SARS-CoV MA15 and measured daily for weight loss. Mice exhibiting extreme weight loss or signs of clinical disease were observed three times a day and euthanized if necessary based on humane endpoints. The virus inoculum dose was selected to result in a range of susceptibility phenotypes in the 8 founder strains. Previous studies were performed on a C57BL/6 background, so this dose was then tested in the founder strains to ensure a range of susceptibility, mortality, and immune responses. We aimed to maximize phenotypic diversity while still maintaining sufficient survival such that we could assess immune phenotypes at various times post-infection.

### Flow cytometry

Spleens were prepared for flow cytometry staining as previously described (Graham et al., 2017a; Graham et al., 2017b; Graham et al., 2016; Graham et al., 2015). All antibodies were tested using cells from the 8 CC founder strains to confirm that antibody clones were compatible with the CC mice prior to being used for testing.

### Statistical analysis

When comparing groups, Mann-Whitney tests were conducted, with p-values <0.05 considered significant. Error bars are +/− SD. Linear regression analysis was performed using GraphPad Prism software.

## Results

### Infection of genetically diverse mice with SARS-CoV results in a variety of viral load trajectories

As part of a screen of genetically diverse mice from the CC for clinical outcomes and immune phenotypes following SARS-CoV MA15 infection, 18-28 mice each from over 100 different CC-RIX lines were infected with SARS-CoV MA15, followed by monitoring for survival and weight loss up to 28 days post-infection. In addition, lung viral loads were measured at days 2 and 4 post-infection using separate cohorts of mice. Infection of CC-RIX mice with SARS-CoV MA15 resulted in wide range of average lung viral loads at 2 days post-infection, ranging from below the limit of detection to 4.75×10^7^ PFU (**Figure 1A**). Furthermore, while the vast majority of CC-RIX lines experienced a decrease in average viral loads from day 2 to day 4 post-infection, the amount of decrease varied considerably (**Figure 1B**). In order to investigate the immune correlates of early viral control upon infection, we examined selected lines with extreme phenotypes for further examination. As shown in Figure 1A, lines with an average lung viral load of less than 10^5^ at day 2 post-infection (N=8) were considered to be “low titer”, and lines with an average lung viral load of greater than 10^7^ at day 2 post-infection (N=24) were considered to be “high titer” for further analysis (**Figure 1C** and **Supplementary Table 1**).

**Figure 1.**
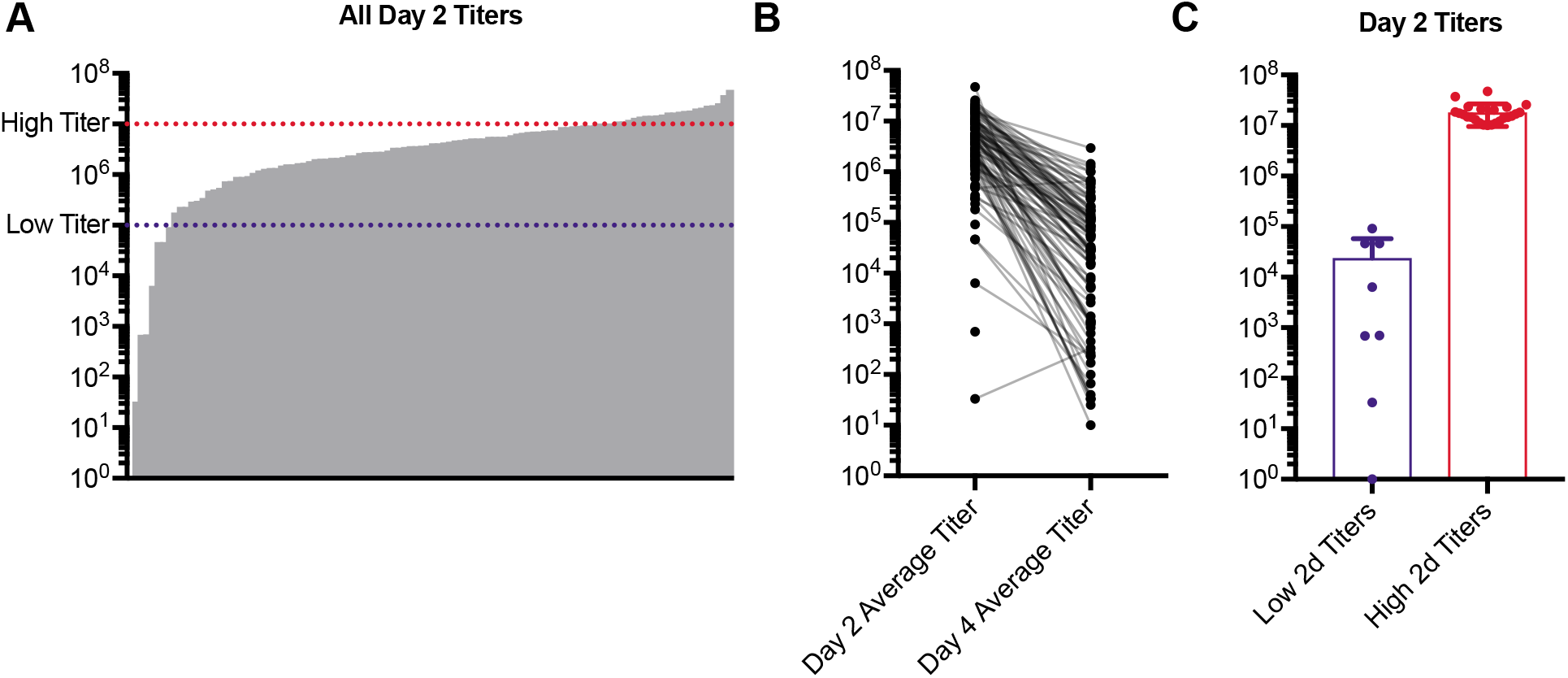
SARS-CoV MA15 infection of genetically diverse mice results in a variety of viral load trajectories. Age-matched female CC-RIX were infected intranasally with SARS-CoV MA15. (A) Average viral loads in the lung at day 2 post-infection are shown for each CC-RIX line. Red dotted line indicates titers above 10^7^ PFU, and blue dotted line indicates viral titers below 10^5^ PFU. (B) Average viral loads in the lung at day 2 post-infection and at day 4 post-infection for each CC-RIX line. (C) The day 2 post-infection average lung viral loads are shown for selected CC-RIX lines are with extreme phenotypes: low or high viral titers. Lines with an average lung viral load of less than 10^5^ at day 2 post-infection (N=8) were considered to be “low titer”, and lines with an average lung viral load of greater than 10^7^ at day 2 post-infection (N=24) were considered to be “high titer” for further analysis.

### Early viral control in the lung correlates with distinct T cell phenotypes and inflammatory potential

In order to determine baseline immune signatures that correlate with progression to high viral load upon infection, we examined the frequency of different populations and phenotypes of T cells within the spleen (as a proxy for the circulation) at steady state by assessment of a second cohort of age-matched mice from each of these CC-RIX lines (**Figure 1C** and **Supplementary Table 1**). CC-RIX mice with superior virologic containment at day 2 post-infection had a higher mean frequency of CD44+ CD4 and CD8 T cells in the spleen prior to infection (**Figures 2A-B**), in addition to an increased proportion of CD4 T cells that express Ki67 (**Figure 2C**), which signals recent proliferation. Along with this increase in the frequency of CD44+ memory T cells, mice from CC-RIX lines with low viral titers at day 2 post-infection had a significantly increased frequency of baseline splenic Foxp3+ regulatory T cells (Treg) (**Figure 2D**). Furthermore, mice from these lines had an increased frequency of Tregs that are CD44+ (**Figure 2E**) and that are CD73+ (**Figure 2F**). In addition to these significant findings, we assessed a variety of activation markers on conventional CD4 and CD8 T cells as well as Tregs at steady state, many of which are not different between the two groups (**Figures 2G-H**). Finally, there is a statistically significant positive correlation between the frequency of regulatory T cells and CD44+ CD4+, CD44+ CD8+, and Ki67+ CD4+ T cells independent of SARS-COV MA15 viral outcomes (**Figures 2I-K**). Together, these data suggest that mice that are better able to contain virus replication early following infection have a higher baseline circulating frequency of both memory T cells as well as regulatory T cells in the spleen.

**Figure 2.**
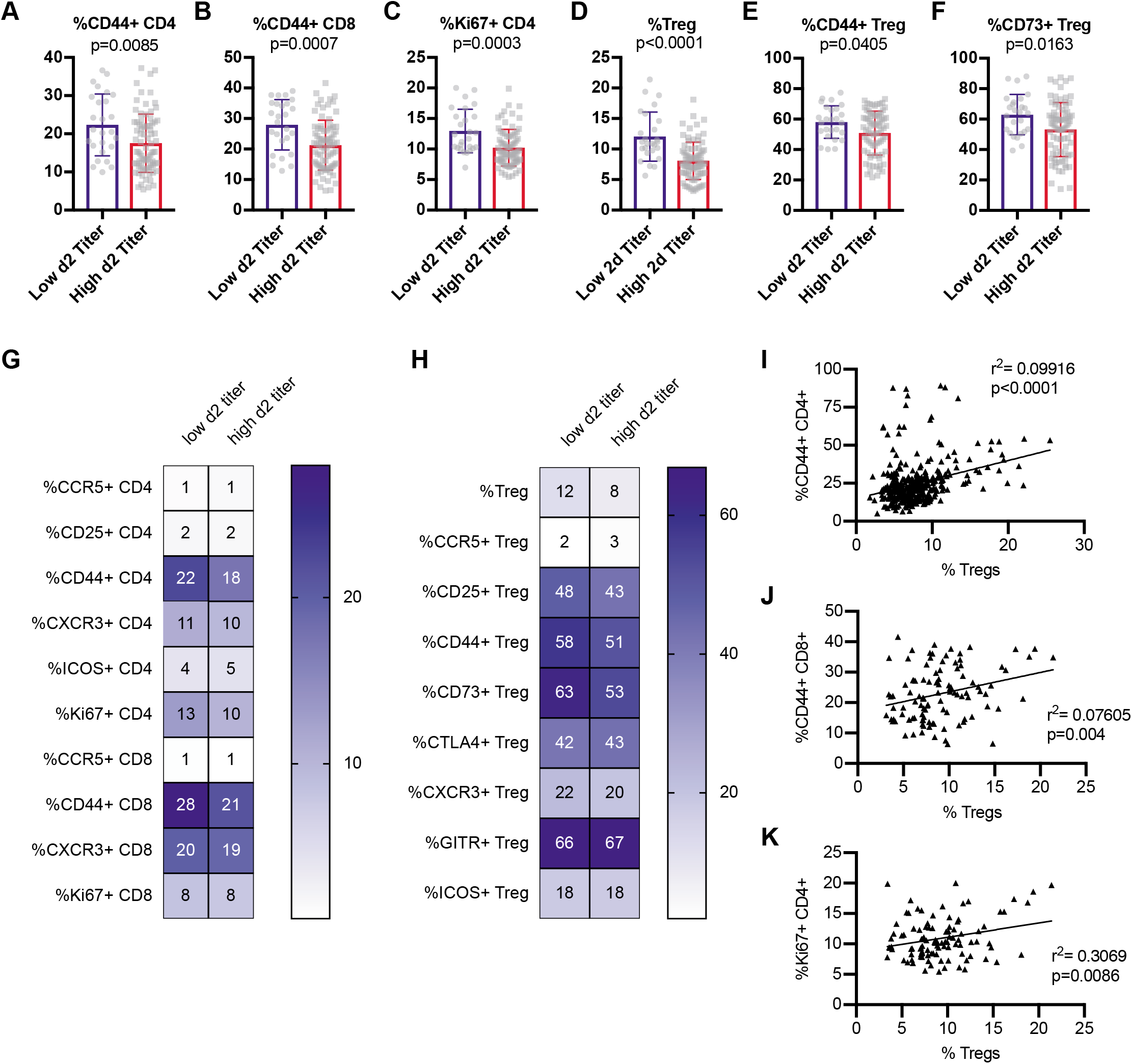
Early virologic control correlates with increased baseline circulating frequency of activated T cells and regulatory T cells. Age-matched female CC-RIX were infected intranasally with SARS-CoV MA15 and lung viral loads at day 2 post-infection were used to select CC-RIX lines with extreme phenotypes: “Low 2d Titer” or “High 2d Titer”, as indicated in Figure 1. Mice from a second cohort of 3-6 age-matched male mice of these selected lines were euthanized and splenic cells analyzed by flow cytometry staining to determine the % of CD4 T cells that are CD44+ (A), the % of CD8 T cells that are CD44+ (B), the % of CD4 T cells that are Ki67+ (C), the % of CD4 T cells that are Foxp3+ Tregs (D), the % of Tregs that are CD44+ (E), and the % of Tregs that are CD73+ (F). Statistical significance was determined by Mann-Whitney test. Heat maps were made to compare the average percent of the indicated cell populations for conventional T cells (G) and for regulatory T cells (H). No statistical significance (p>0.05 by Mann-Whitney test) was found for any comparisons except those indicated in Figures 2A-F. The correlation between the baseline splenic frequency of Tregs (% Foxp3+ of CD4 T cells) and (I) % of CD4 T cells that are CD44+, (J) % of CD8 T cells that are CD44+, or (K) % of CD4 T cells that are Ki67+ are shown with linear regressions for mice from all CC-RIX lines with low or high day 2 titer.

Next, we assessed the ability of T cells to express cytokines at steady state by stimulating baseline splenocytes polyclonally using an *ex vivo* intracellular cytokine stimulation assay. Mice from CC-RIX lines that had a low lung viral titer at day 2 post-infection had an increased frequency of baseline splenic CD8 T cells that could express IFNg (**Figure 3A**) as well as IL-17 (**Figure 3B**). Additionally, an increased frequency of steady-state splenic CD4 T cells that express IL-17 upon polyclonal stimulation was found in mice from CC-RIX lines with low lung viral loads at day 2 post-infection (**Figure 3C**). Upon examination of T cells expressing a combination of TNFa and IFNg, we found that mice from lines with superior early virologic control had an increased frequency of CD8 T cells that were TNFa-IFNg+ (**Figure 3D**) and a decreased frequency that were TNFa+IFNg- (**Figure 3E**). Similarly, mice from lines with high viral titers at day 2 post-infection had an increased fraction of baseline circulating CD4 T cells that express TNFa (**Figure 3F**), as well as an increased fraction of CD4 T cells that are TNFa+IFNg- (**Figure 3G**). Taken together, our results suggest that early viral control upon infection with SARS-CoV MA15 correlates with a pre-infection increased frequency of circulating T cells with a potential to express IFNg or IL17 rather than TNFa (**Figure 3H**). This latter finding is consistent with previous studies of SARS-CoV that found TNFa to play a detrimental role in tissue damage after infection (McDermott et al., 2016), and therefore may serve as a biomarker for individuals who may be at higher risk of high viral loads upon CoV infections. Notably, there is a significant positive correlation between the frequency of splenic Tregs at baseline and the expression of IL-17 by CD4 or CD8 T cells, and a negative correlation between baseline frequency of Tregs in the spleen and TNFa expression by CD4 or CD8 T cells (**Figure 3I**), further underscoring the potential immunoprotective signature linked with baseline Treg frequency.

**Figure 3.**
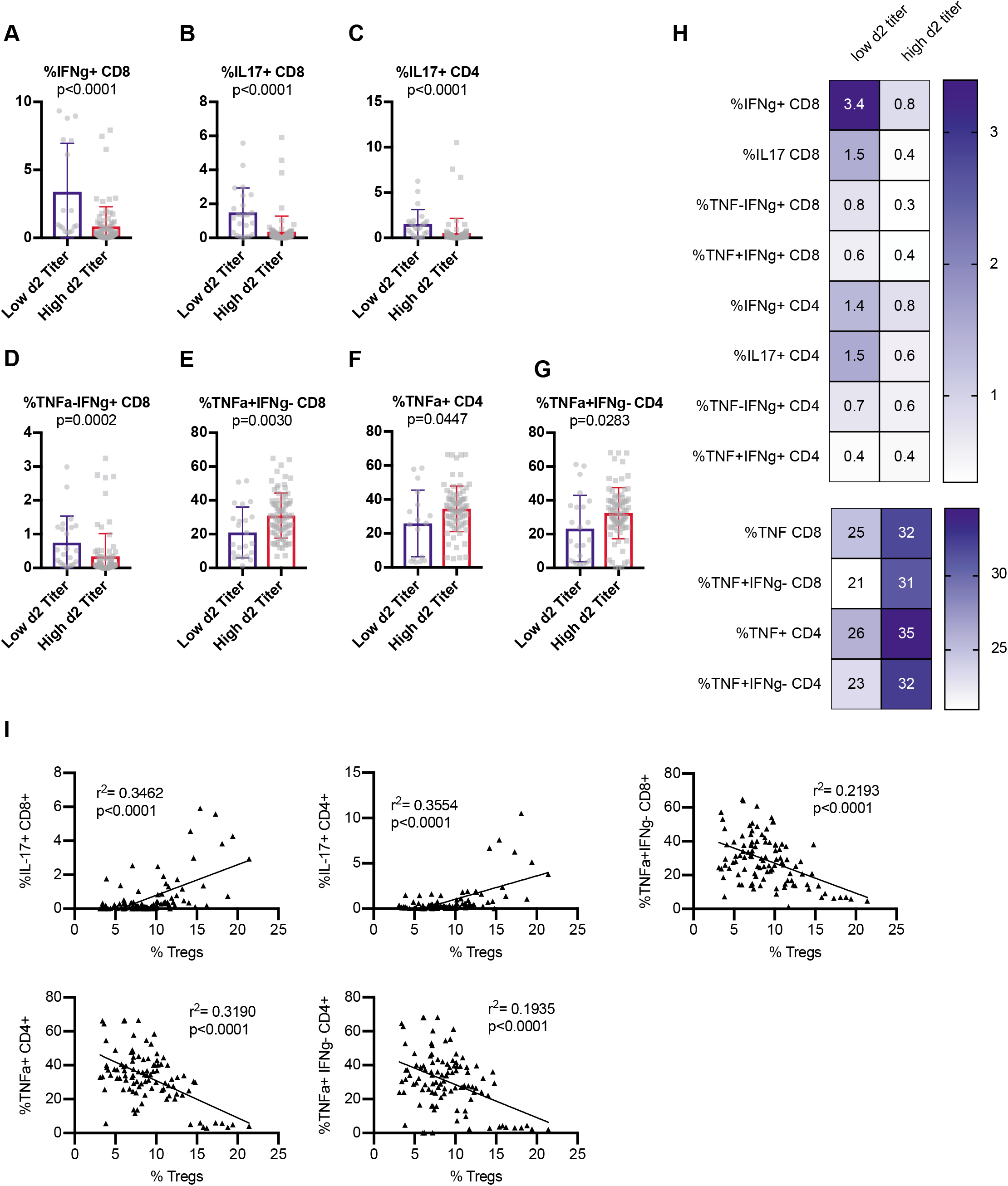
Early viral control upon infection correlates with baseline T cells with a potential to express IFNg or IL17 rather than TNF. Age-matched female CC-RIX were infected intranasally with SARS-CoV MA15 and lung viral loads at day 2 post-infection were used to select CC-RIX lines with extreme phenotypes: “Low 2d Titer” or “High 2d Titer”, as indicated in Figure 1. Mice from a second cohort of 3-6 age-matched male mice of these selected lines were euthanized and splenic cells were treated with anti-CD3/CD28 for intracellular cytokine staining assessment of (A) %IFNg+ of CD8 T cells, (B) %IL-17+ of CD8 T cells, (C) %IL-17+ of CD4 T cells, (D) %TNF-IFNg+ of CD8 T cells, (E) %TNF+IFNg-of CD8 T cells, (F) %TNF+ of CD4 T cells, and (G) %TNF+IFNg-of CD4 T cells. Statistical significance was determined by Mann-Whitney test. (H) Heat maps were made to compare the average percent of the indicated cell populations. No statistical significance (p>0.05 by Mann-Whitney test) was found for any comparisons except those indicated in Figures 3A-G. (I) The correlation between the baseline splenic frequency of Tregs (% Foxp3+ of CD4 T cells) and % of CD8 T cells that are IL-17+, % of CD4 T cells that are IL-17+, % of CD8 T cells that are TNF+IFNg-, % of CD4 T cells that are TNF+, and the % of CD4 T cells that are TNF+IFNg-are shown with linear regressions for mice from all CC-RIX lines with low or high day 2 titer.

### Circulating T cell phenotypes at steady state predict protection from high titers and disease upon SARS-CoV MA15 infection

To identify possible baseline immune predictors of both severe virologic and disease outcomes upon infection, we classified CC-RIX lines with extreme phenotypes based on both lung viral loads at days 2 and 4 post-infection, as well as weight loss and mortality. Lines were categorized as “low infection and disease” (LID), which had 0-5% weight loss upon infection, no death, day 2 average lung viral titers of <10^5^ and average day 4 lung viral titers of <10^4^ (N=5 lines). Conversely, N=4 lines were categorized as “high infection and disease” (HID) if they experienced greater than 15% weight loss and death, as well as average lung viral titers at day 2 post-infection of >10^6^ and average lung viral titers at day 4 post-infection of >10^5^ (**Figure 4A** and **Supplemental Table 1**). Upon examination of splenic baseline T cell phenotypes in mice from these 9 CC-RIX lines, we found a significantly elevated CD4:CD8 T cell ratio in mice from lines that experienced low infection and disease compared to those that had high infection and disease (**Figure 4B**). Similar to what we found when considering day 2 post-infection viral titers alone, we found that a higher frequency of circulating CD44+ CD8 T cells at baseline correlated with protection from high infection and disease (**Figure 4C**), whereas a lower frequency of CCR5+ or CD25+ CD4 T cells correlated with protection from high viral loads and disease (**Figures 4D-E**). In addition to conventional T cells, we also assessed the ability of circulating Treg frequency and phenotype to predict viral load and disease outcomes upon SARS-CoV MA15 infection. An increased baseline frequency of circulating Tregs was present in mice from LID CC-RIX lines (**Figure 4F**). Mice from CC-RIX lines with low infection and disease had a reduced frequency of Tregs expressing CD25 or CCR5 (**Figures 4G-H**), but an increased frequency of Tregs expressing CD73 (**Figure 4I**). Thus, it is possible that Treg migration patterns and/or mechanisms of suppression may influence the virologic and clinical outcomes upon SARS-CoV infection.

**Figure 4.**
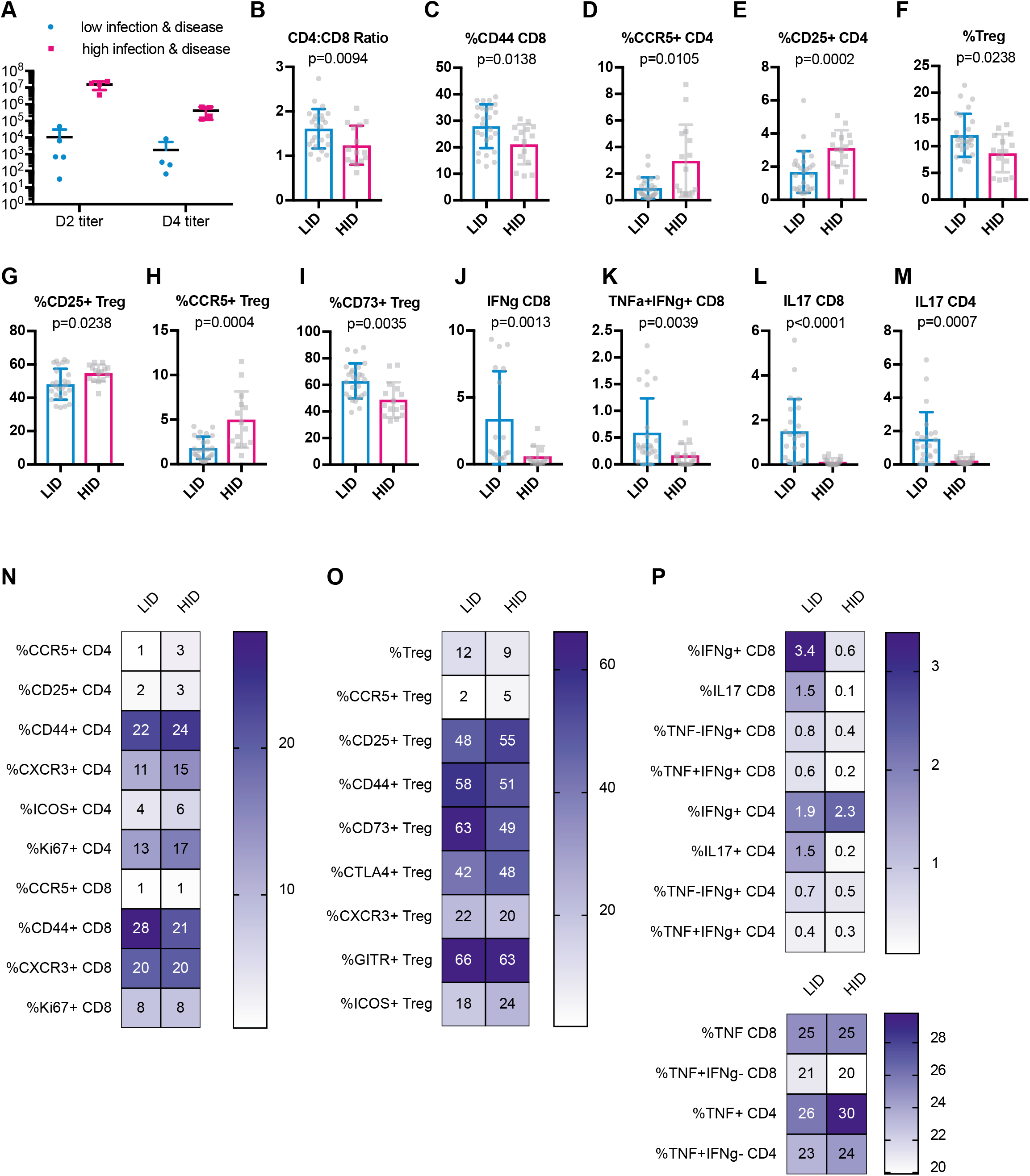
Baseline activated CD8 T cells and Tregs correlate with severe virologic and disease outcomes upon SARS-CoV infection. Age-matched female CC-RIX were infected intranasally with SARS-CoV MA15 and mice were monitored for death, weight loss, and lung viral loads. To identify possible baseline immune predictors of both viral replication as well as disease upon infection, we classified CC-RIX lines with extreme phenotypes based on both lung viral loads at days 2 and 4 post-infection, as well as weight loss and mortality. Lines were categorized as “low infection and disease” (LID), which had 0-5% weight loss upon infection, no death, day 2 average lung viral titers of <10^5^ and average day 4 lung viral titers of <10^4^ (N=5 lines). Conversely, N=4 lines were categorized as “high infection and disease” (HID) if they experienced greater than 15% weight loss and death, as well as average lung viral titers at day 2 post-infection of >10^6^ and average lung viral titers at day 4 post-infection of >10^5^. Lung viral titers from these 9 CC-RIX lines are shown for days 2 and 4 post-infection (A). Mice from a second cohort of 3-6 age-matched male mice of these selected 9 lines were euthanized and splenic cells analyzed by flow cytometry staining to determine the CD4:CD8 ratio (B), % of CD8 T cells that are CD44+ (C), % of CD4 T cells that are CCR5+ (D), % of CD4 T cells that are CD25+ (E), % of CD4 T cells that are Foxp3+ Treg (F), % of Tregs that are CD25+ (G), % of Tregs that are CCR5+ (H), and % of Tregs that are CD73+ (I). In addition, splenic cells were treated with anti-CD3/CD28 for intracellular cytokine staining assessment of (J) %IFNg+ of CD8 T cells, (K) %TNF+IFNg+ of CD8 T cells, (L) %IL-18+ of CD8 T cells, and (M) %IL-17+ of CD4 T cells. Statistical significance was determined by Mann-Whitney test. (N-P) Heat maps were made to compare the average percent of the indicated cell populations. No statistical significance (p>0.05 by Mann-Whitney test) was found for any comparisons except those indicated in Figures 4B-M.

Finally, we assessed the potential of T cells to express cytokines at baseline. Mice from CC-RIX lines with low infection and disease had increased expression of IFNg upon polyclonal *ex vivo* stimulation (**Figure 4J**), as well as increased co-expression of both IFNg and TNFa (**Figure 4K**). Additionally, mice from CC-RIX lines with low lung viral loads and low disease upon infection also had an increased circulating fraction of splenic CD8 and CD4 T cells that express IL-17 upon stimulation (**Figures 4L-M**). Altogether, our findings suggest a distinct circulating T cell signature at steady-state that is associated with severe virologic and clinical outcomes upon SARS-CoV infection (**Figures 4O-P**).

### Dysregulated circulating T cell phenotypes at steady state are associated with disease in the setting of high viral loads upon SARS-CoV MA15 infection

To further improve our understanding of why some individuals experience severe illness and disease upon infection while others do not, we wished to further investigate immune correlates of protection from disease when viral loads were normalized. Thus, to identify possible baseline immune predictors of disease upon infection with a high early lung viral load, we differently classified CC-RIX lines with extreme phenotypes based on both lung viral loads at days 2 and 4 post-infection, as well as weight loss and mortality. Lines were categorized as “no disease high titer” (NDHT), which had 0-5% weight loss upon infection and no death despite day 2 average lung viral titers of >10^7^ and average day 4 lung viral titers of >10^5^ (N=3 lines) and “disease high titer” (DHT; N=3 lines) if they experienced greater than 15% weight loss and death, as well as average lung viral titers at day 2 post-infection of >10^7^ and average lung viral titers at day 4 post-infection of >10^5^ (Supplemental Table 1). Thus, there were no differences in average viral loads between groups (Figure 5A), and we could assess how baseline T cell phenotypes correlated with eventual disease upon similar levels of infection. We found that there was a significantly elevated CD4:CD8 T cell ratio in mice from lines that experienced no disease in the context of high viral loads compared to those that showed signs of disease (Figure 5B). However, upon examination of the phenotype of these CD4 T cells, we found that a decreased baseline frequency of CD25+ or CCR5+ circulating CD4 T cells was associated with protection from disease (Figure 5C-D). In addition, mice from CC-RIX lines that were protected from disease in a setting of high viral loads had a reduced fraction of Tregs that expressed CD25 or CTLA-4 (Figures 5E-F). Finally, mice from lines that were protected from disease had baseline circulating CD8 T cells that were more likely to express both TNFa and IFNg upon polyclonal stimulation (Figure 5G), thereby indicating that this could be a predictor of protection from disease upon infection. In sum, our findings suggest a baseline circulating signature of T cell dysfunction is associated with severe clinical outcomes upon SARS-CoV infection with high levels of early virus replication (Figures 5H-J).

**Figure 5.**
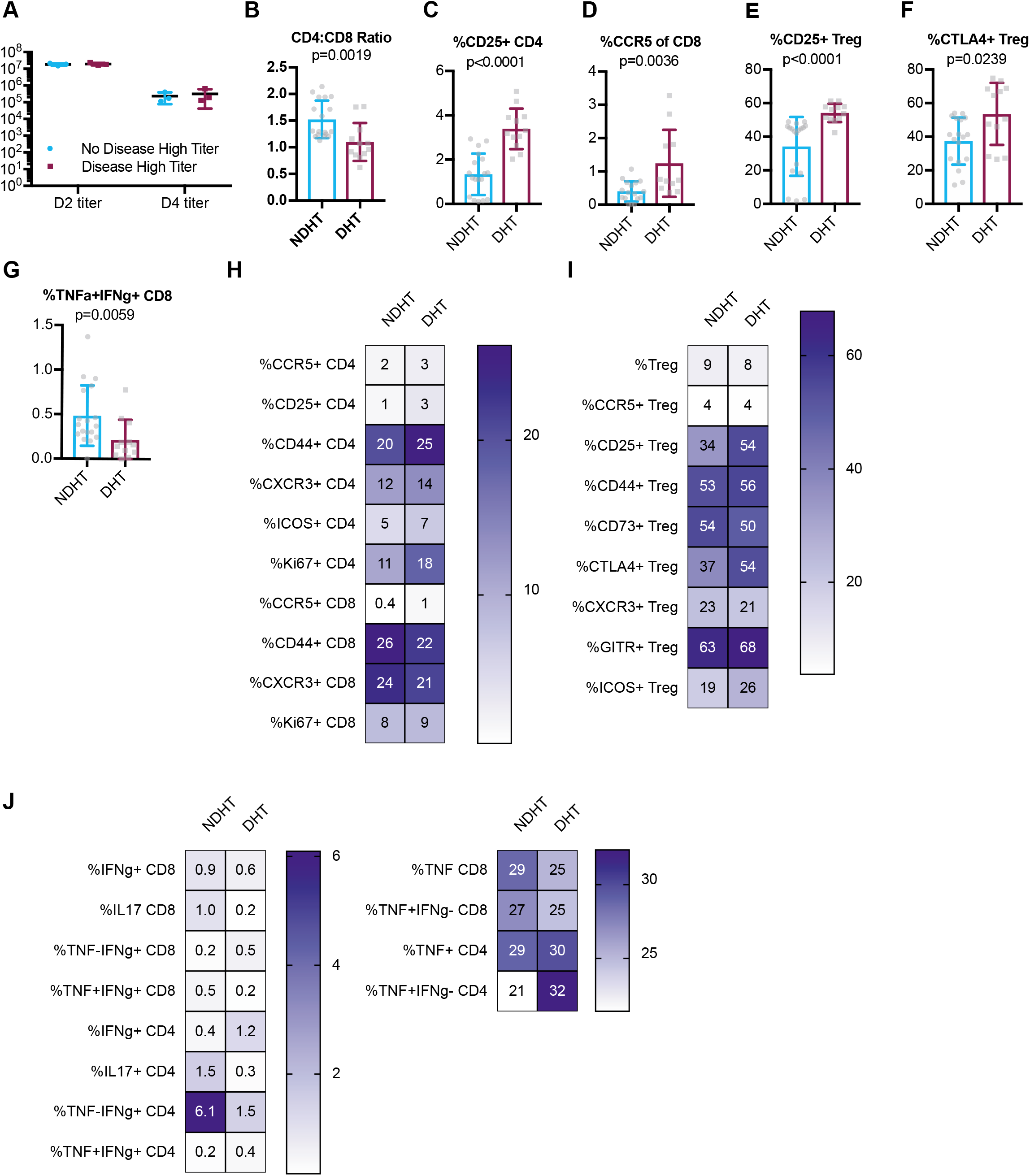
A dysregulated circulating baseline T cell phenotype is associated with severe disease in the setting of high viral loads upon infection. Age-matched female CC-RIX were infected intranasally with SARS-CoV MA15 and mice were monitored for death, weight loss, and lung viral loads. To identify possible baseline immune predictors of disease upon infection with a high early lung viral load, we classified CC-RIX lines with extreme phenotypes based on both lung viral loads at days 2 and 4 post-infection, as well as weight loss and mortality. Lines were categorized as “no disease high titer” (NDHT), which had 0-5% weight loss upon infection and no death despite day 2 average lung viral titers of >10^7^ and average day 4 lung viral titers of >10^5^ (N=3 lines) and “disease high titer” (DHT; N=3 lines) if they experienced greater than 15% weight loss and death, as well as average lung viral titers at day 2 post-infection of >10^7^ and average lung viral titers at day 4 post-infection of >10^5^. Lung viral titers from these 6 CC-RIX lines are shown for days 2 and 4 post-infection (A). Mice from a second cohort of 3-6 age-matched male mice of these selected 6 lines were euthanized and splenic cells analyzed by flow cytometry staining to determine the CD4:CD8 ratio (B), % of CD4 T cells that are CD25+ (C), % of CD8 T cells that are CCR5+ (D), % of Tregs that are CD25+ (E), and % of Tregs that are CTLA-4+ (F). In addition, splenic cells were treated with anti-CD3/CD28 for intracellular cytokine staining assessment of (G) %TNF+IFNg+ of CD8 T cells. Statistical significance was determined by Mann-Whitney test. (H-J) Heat maps were made to compare the average percent of the indicated cell populations. No statistical significance (p>0.05 by Mann-Whitney test) was found for any comparisons except those indicated in Figures 5B-G.

## Discussion

The COVID-19 pandemic poses enormous challenges to global healthcare systems, as healthcare workers struggle to find adequate personal protective equipment (PPE) with which to shield themselves as they attempt to treat patients with a single FDA-authorized drug for emergency use, remdesivir (Pruijssers et al., 2020; Sheahan et al., 2017), and without access to a protective vaccine. While the latter are under rapid development, it is clear that as new treatment and prevention strategies are developed, there will likely be an inadequate supply available for all those in need. This underscores the need to be able to identify individuals at highest risk of infection and disease to be able to best triage protective PPE as well as newly developed treatment and prevention strategies, including vaccines. Further, the concept of “super-spreaders”, or rare individuals with a unique capacity to infect a large number of individuals (Goyal et al., 2020), suggests that virologic control and identification of individuals who may be most prone to high viral loads may be critical to limit and/or halt the spread of SARS-CoV-2. While many immune correlates of severe disease upon infection with SARS-CoV-2 have been recently identified in humans, to date these studies involve analysis of already infected individuals who present with mild or severe illness, as compared to healthy controls. Therefore, it is difficult to determine whether immune signatures from these individuals are predictive, or rather represent symptoms associated with specific disease states.

In the absence of prospectively collected, pre-SARS-CoV-2 infection human samples that could be used for a case-control analysis to allow for identification of predictive immune signatures of COVID-19 virologic and clinical outcomes, we utilized a mouse model system to identify baseline, circulating T cell signatures that predict severe infection and disease outcomes upon SARS-CoV infection. Use of the CC mouse model population enabled the study of a diversity of virologic and disease outcomes upon infection with SARS-CoV, as the genetic diversity inherent to the model better replicates the genetic diversity in the human population, and thus contributes to diverse phenotypes, including immunophenotypes and disease phenotypes pre- and post-infection. The use of the mouse-adapted SARS-CoV MA15, while not the same as SARS-CoV-2, at the very least allowed us to perform proof-of-concept studies demonstrating that baseline T cell phenotypes can predict infection and disease outcomes following coronavirus infections, though future studies of both mice as well as human samples using SARS-CoV-2 are required to validate our findings for COVID-19.

In our previous study, we used data from our screen of CC mice to identify more universal immune correlates of mortality following infection with influenza, SARS-CoV, and WNV (Graham et al., 2020). The protective signature included an increased level of basal T-cell activation that was associated with protection, which we also found here to be associated with protection from severe virologic and clinical outcomes following SARS-CoV infection (**Figures 2, 4, & 5**). As CD44 is a T cell marker associated with antigen experience or a memory phenotype, it is possible that these memory T cells could undergo rapid bystander activation via the innate immune response following CoV infection, and thus play a critical early role in a protective response. The presence of these CD44+ T cells may indicate true, conventional memory T cells that resulted from previous microbial exposures in the murine specific pathogen-free (SPF) colony. Alternatively, such cells could also be unconventional memory cells that possess a memory phenotype despite not having encountered cognate antigen (Jameson, 2005; Le Campion et al., 2002; Min et al., 2003; Schuler et al., 2004; Surh and Sprent, 2005). Nevertheless, CD44+ T cells of either origin could participate in early viral control through bystander-mediated activity and thus confer a protective advantage through rapid viral containment before the virus-specific T cell response has been generated (Chu et al., 2013). Such activity is consistent with previous work demonstrating that unconventional memory T cells can aid in protection against pathogens including *Listeria monocytogenes* and influenza virus (Lanzer et al., 2018; Lee et al., 2013; Sosinowski et al., 2013).

It stands to reason that such an active innate-like T cell response would need to be subject to immunoregulation in order to limit activity and prevent excess collateral damage. Also in our previous study, we found that an increased frequency of Tregs correlated with protection from death following each of the three infections (influenza A virus, West Nile virus, and SARS-CoV) (Graham et al., 2020). Our results presented herein further support that an increased basal frequency of Tregs in the circulation correlates with protection both from early SARS-CoV viral replication, as well as from disease upon infection (**Figures 2 and 4**). In the context of multiple viral infections, we and others have found that Tregs are critical to orchestrate proper anti-viral immune responses (Lanteri et al., 2009; Lund et al., 2008; Pattacini et al., 2016; Ruckwardt et al., 2009; Soerens et al., 2016), while it has also been found that Tregs in the context of infections, including respiratory infections such as RSV and influenza, can assist in protecting the host from excessive immunopathology (Belkaid and Tarbell, 2009; Brincks et al., 2013; Lee et al., 2010; Loebbermann et al., 2012; Richert-Spuhler and Lund, 2015; Smigiel et al., 2014). Thus, our results here further support the concept that balance between anti-viral immunity and immunoregulation is essential to spare the host from both unrestricted viral replication as well as severe disease after infection. We predict that Tregs play this dual role in the context SARS-CoV infection as well, wherein their increased abundance at steady state (**Figures 2D and 4F**) is advantageous in terms of allowing for the generation of an appropriately focused anti-viral immune response, while variable expression of particular homing and activation markers allows for an appropriately tuned suppressive response. While a complete characterization of Tregs after infection would help to reveal the dynamics of an appropriate Treg response in the context of SARS-CoV infection, we do not have this data from our screen, and so further studies are needed to fully assess Treg phenotype and function in both mice and humans after SARS-CoV and SARS-CoV-2.

Finally, in both our previous study as well as this focused study of SARS-CoV, we found that a restricted pro-inflammatory potential of T cells is correlated with protection from mortality upon infection with each of the three viruses (Graham et al., 2020) as well as severe virologic outcomes upon SARS-CoV infection (**Figures 3-5**). Specifically, we demonstrate that pre-infection ability of T cells to express the pro-inflammatory cytokine TNF correlated with more severe virologic outcomes (**Figures 3E-G**), as has been demonstrated as well for SARS-CoV and COVID-19 (Blanco-Melo et al., 2020; Lucas et al., 2020; McDermott et al., 2016; Qin et al., 2020). On the other hand, the presence of circulating T cells at steady-state with the potential to express IFNg or IL-17 is associated with protection from both early and high lung viral loads (**Figures 3A-D and 4J-M**) as well as disease (**Figures 4J-M and Figure 5G**). IFNg is well known as a potent anti-viral cytokine, and so it is not a surprise that this cytokine could play a role in SARS-CoV restriction, and though the potential role of IL-17 is less clear.

Overall, the results from our study demonstrate that baseline T cell phenotypes can predict early virologic and clinical outcomes upon infection with SARS-coronaviruses. While it is clear that additional mechanistic and human studies are needed to validate these findings for extrapolation to COVID-19, this study also serves to highlight the complexity of inflammation, which can at the same time be protective and detrimental to the host. We hypothesize that particular T cell immunophenotypes or signatures may be critical to promoting rapid immunity upon infection and limiting immune-mediated collateral damage, and further predict that bystander-activated T cells may play a powerful role in the early innate immune response to SARS-CoV. However, as COVID-19 is associated with more inflammatory responses than SARS, the correlates of disease and protection for SARS-CoV-2 may differ from those of SARS-CoV. Thus, future studies include using select CC strains with extreme baseline immune phenotypes to validate our findings with SARS-CoV MA15 as well as mouse-adapted SARS-CoV-2 (Dinnon et al., 2020). Alternatively, usage of transient depletion systems, such as the Foxp3^DTR^ mouse model (Kim et al., 2007), would enable targeted elimination of all or some Foxp3+ Tregs prior to infection with SARS-CoV or SARS-CoV-2 in order to directly test the role of Tregs in SARS-CoV virologic and clinical outcomes. Nevertheless, our data presented herein support the concept that levels of inflammation prior to coronavirus infection may impact post-infection virologic and clinical disease states.

## Acknowledgements

We wish to thank our collaborators in the Systems Immunogenetics Group for helpful discussions and generation of mice. In particular, we wish to thank Ginger Shaw for her tireless work generating the RIX mice used in this study. Funding for this study was provided by NIH grant U19AI100625.

## Author Contributions

DRM, SKM, MTF, FPMV, MTH, RSB, and JML designed the research studies; JBG, JLS, SRL, VDM, LEG, and AS conducted experiments and acquired and analyzed data; SJ and MAM performed data cleaning and integration; and JBG and JML wrote the first draft of the manuscript. All authors read the manuscript and contributed editorial suggestions.

The authors have declared that no conflict of interest exists.

Funding for this study was provided by NIH grant U19AI100625.

---

**Supplemental Table 1.** CC F1 lines in infection and disease categories

| **Supplemental Table 1. CC F1 lines in infection and disease categories** |  |  |  |
| --- | --- | --- | --- |
| **RIX Line** | **d2 Viral Load** | **D4 Viral Load** | **Group** |
| CC042xCC019 | 0 | 847 | Low titer |
| CC030xCC061 | 33 | 333 | Low titer |
| CC018xCC065 | 683 | N/A | Low titer |
| CC030xCC023 | 700 | 0 | Low titer |
| CC052xCC014 | 6333 | 233 | Low titer |
| CC012xCC032 | 46333 | 233 | Low titer |
| CC032xCC013 | 46667 | 67 | Low titer |
| CC011xCC032 | 91500 | 0 | Low titer |
| CC019xCC004 | 10200000 | 27000 | High titer |
| CC068xCC043 | 10350000 | 105750 | High titer |
| CC006xCC039 | 10366667 | 211000 | High titer |
| CC057xCC052 | 11193333 | 28000 | High titer |
| CC062xCC046 | 11333333 | 114667 | High titer |
| CC029xCC071 | 12166667 | 577500 | High titer |
| CC060xCC037 | 13200000 | 14470 | High titer |
| CC041xCC016 | 14100000 | 120700 | High titer |
| CC028xCC024 | 14650000 | 2966667 | High titer |
| CC065xCC010 | 14683333 | 330667 | High titer |
| CC001xCC055 | 14800000 | N/A | High titer |
| CC013xCC041 | 15366667 | 172500 | High titer |
| CC074xCC058 | 16800000 | 128667 | High titer |
| CC004xCC011 | 16816667 | 1003333 | High titer |
| CC026xCC034 | 17500000 | 175667 | High titer |
| CC061xCC025 | 18200000 | 192667 | High titer |
| CC016xCC038 | 18666667 | 116000 | High titer |
| CC056xCC033 | 20333333 | 300033 | High titer |
| CC033xCC046 | 21000000 | 409667 | High titer |
| CC025xCC028 | 23166667 | 640333 | High titer |
| CC042xCC025 | 23666667 | N/A | High titer |
| CC015xCC059 | 26000000 | 80000 | High titer |
| CC055xCC006 | 37333367 | N/A | High titer |
| CC055xCC028 | 47566667 | 33 | High titer |
| CC018xCC065 | 683 | 8400 | LID |
| CC032xCC013 | 46667 | 67 | LID |
| CC030xCC023 | 700 | 0 | LID |
| CC030xCC023 | 33 | 333 | LID |
| CC052xCC014 | 6333 | 233 | LID |
| CC001xCC074 | 3686667 | 688833 | HID |
| CC074xCC058 | 16800000 | 128667 | HID |
| CC025xCC028 | 23166667 | 640333 | HID |
| CC061xCC025 | 18200000 | 192667 | HID |
| CC013xCC041 | 15366667 | 172500 | NDHT |
| CC016xCC038 | 18666667 | 116000 | NDHT |
| CC033xCC046 | 21000000 | 409667 | NDHT |
| CC074xCC058 | 16800000 | 128667 | DHT |
| CC025xCC028 | 23166667 | 640333 | DHT |
| CC061xCC025 | 18200000 | 192667 | DHT |

## References

Belkaid, Y., and K. Tarbell. 2009. Regulatory T cells in the control of host-microorganism interactions (*). Annu Rev Immunol 27:551–589.

Blanco-Melo, D., B.E. Nilsson-Payant, W.C. Liu, S. Uhl, D. Hoagland, R. Moller, T.X. Jordan, K. Oishi, M. Panis, D. Sachs, T.T. Wang, R.E. Schwartz, J.K. Lim, R.A. Albrecht, and B. R. tenOever. 2020. Imbalanced Host Response to SARS-CoV-2 Drives Development of COVID-19. Cell 181:1036–1045 e1039.

Brincks, E.L., A.D. Roberts, T. Cookenham, S. Sell, J.E. Kohlmeier, M.A. Blackman, and D.L. Woodland. 2013. Antigen-specific memory regulatory CD4+Foxp3+ T cells control memory responses to influenza virus infection. Journal of immunology 190:3438–3446.

Brinkmeyer-Langford, C.L., R. Rech, K. Amstalden, K.J. Kochan, A.E. Hillhouse, C. Young, C. J. Welsh, and D.W. Threadgill. 2017. Host genetic background influences diverse neurological responses to viral infection in mice. Sci Rep 7:12194.

Chu, T., A.J. Tyznik, S. Roepke, A.M. Berkley, A. Woodward-Davis, L. Pattacini, M.J. Bevan, D. Zehn, and M. Prlic. 2013. Bystander-activated memory CD8 T cells control early pathogen load in an innate-like, NKG2D-dependent manner. Cell Rep 3:701–708.

Churchill, G.A., D.C. Airey, H. Allayee, J.M. Angel, A.D. Attie, J. Beatty, W.D. Beavis, J.K. Belknap, B. Bennett, W. Berrettini, A. Bleich, M. Bogue, K.W. Broman, K.J. Buck, E. Buckler, M. Burmeister, E.J. Chesler, J.M. Cheverud, S. Clapcote, M.N. Cook, R.D. Cox, J.C. Crabbe, W.E. Crusio, A. Darvasi, C.F. Deschepper, R.W. Doerge, C.R. Farber, J. Forejt, D. Gaile, S.J. Garlow, H. Geiger, H. Gershenfeld, T. Gordon, J. Gu, W. Gu, G. de Haan, N.L. Hayes, C. Heller, H. Himmelbauer, R. Hitzemann, K. Hunter, H.C. Hsu, F.A. Iraqi, B. Ivandic, H.J. Jacob, R.C. Jansen, K.J. Jepsen, D.K. Johnson, T.E. Johnson, G. Kempermann, C. Kendziorski, M. Kotb, R.F. Kooy, B. Llamas, F. Lammert, J.M. Lassalle, P.R. Lowenstein, L. Lu, A. Lusis, K.F. Manly, R. Marcucio, D. Matthews, J.F. Medrano, D.R. Miller, G. Mittleman, B.A. Mock, J.S. Mogil, X. Montagutelli, G. Morahan, D.G. Morris, R. Mott, J.H. Nadeau, H. Nagase, R.S. Nowakowski, B.F. O’Hara, A.V. Osadchuk, G.P. Page, B. Paigen, K. Paigen, A.A. Palmer, H.J. Pan, L. Peltonen-Palotie, J. Peirce, D. Pomp, M. Pravenec, D.R. Prows, Z. Qi, R.H. Reeves, J. Roder, G.D. Rosen, E.E. Schadt, L.C. Schalkwyk, Z. Seltzer, K. Shimomura, S. Shou, M.J. Sillanpaa, L.D. Siracusa, H.W. Snoeck, J.L. Spearow, K. Svenson, L.M. Tarantino, D. Threadgill, L.A. Toth, W. Valdar, F.P. de Villena, C. Warden, S. Whatley, R.W. Williams, T. Wiltshire, N. Yi, D. Zhang, M. Zhang, F. Zou, and C. Complex Trait. 2004. The Collaborative Cross, a community resource for the genetic analysis of complex traits. Nat Genet 36:1133–1137.

Collaborative Cross, C. 2012. The genome architecture of the Collaborative Cross mouse genetic reference population. Genetics 190:389–401.

Dinnon, K.H., 3rd, S.R. Leist, A. Schafer, C.E. Edwards, D.R. Martinez, S.A. Montgomery, A. West, B.L. Yount, Jr., Y.J. Hou, L.E. Adams, K.L. Gully, A.J. Brown, E. Huang, M.D. Bryant, I.C. Choong, J.S. Glenn, L.E. Gralinski, T.P. Sheahan, and R.S. Baric. 2020. A mouse-adapted model of SARS-CoV-2 to test COVID-19 countermeasures. Nature

Dong, E., H. Du, and L. Gardner. 2020. An interactive web-based dashboard to track COVID-19 in real time. Lancet Infect Dis 20:533–534.

Elbahesh, H., and K. Schughart. 2016. Genetically diverse CC-founder mouse strains replicate the human influenza gene expression signature. Sci Rep 6:26437.

Ferris, M.T., D.L. Aylor, D. Bottomly, A.C. Whitmore, L.D. Aicher, T.A. Bell, B. Bradel-Tretheway, J.T. Bryan, R.J. Buus, L.E. Gralinski, B.L. Haagmans, L. McMillan, D.R. Miller, E. Rosenzweig, W. Valdar, J. Wang, G.A. Churchill, D.W. Threadgill, S.K. McWeeney, M.G. Katze, F. Pardo-Manuel de Villena, R.S. Baric, and M.T. Heise. 2013. Modeling host genetic regulation of influenza pathogenesis in the collaborative cross. PLoS Pathog 9:e1003196.

Goyal, A., D.B. Reeves, E.F. Cardozo-Ojeda, J.T. Schiffer, and B.T. Mayer. 2020. Wrong person, place and time: viral load and contact network structure predict SARS-CoV-2 transmission and super-spreading events. medRxiv 2020.2008.2007.20169920.

Graham, J.B., J.L. Swarts, and J.M. Lund. 2017a. A Mouse Model of West Nile Virus Infection. Curr Protoc Mouse Biol 7:221–235.

Graham, J.B., J.L. Swarts, V.D. Menachery, L.E. Gralinski, A. Schafer, K.S. Plante, C.R. Morrison, K.M. Voss, R. Green, G. Choonoo, S. Jeng, D.R. Miller, M.A. Mooney, S.K. McWeeney, M.T. Ferris, F. Pardo-Manuel de Villena, M. Gale, M.T. Heise, R.S. Baric, and J.M. Lund. 2020. Immune Predictors of Mortality After Ribonucleic Acid Virus Infection. J Infect Dis 221:882–889.

Graham, J.B., J.L. Swarts, M. Mooney, G. Choonoo, S. Jeng, D.R. Miller, M.T. Ferris, S. McWeeney, and J.M. Lund. 2017b. Extensive Homeostatic T Cell Phenotypic Variation within the Collaborative Cross. Cell Rep 21:2313–2325.

Graham, J.B., J.L. Swarts, S. Thomas, K.M. Voss, A. Sekine, R. Green, R.C. Ireton, M. Gale, Jr., and J.M. Lund. 2018. Immune correlates of protection from West Nile virus neuroinvasion and disease. J Infect Dis

Graham, J.B., J.L. Swarts, C. Wilkins, S. Thomas, R. Green, A. Sekine, K.M. Voss, R.C. Ireton, M. Mooney, G. Choonoo, D.R. Miller, P.M. Treuting, F. Pardo Manuel de Villena, M.T. Ferris, S. McWeeney, M. Gale, Jr., and J.M. Lund. 2016. A Mouse Model of Chronic West Nile Virus Disease. PLoS Pathog 12:e1005996.

Graham, J.B., S. Thomas, J. Swarts, A.A. McMillan, M.T. Ferris, M.S. Suthar, P.M. Treuting, R. Ireton, M. Gale, Jr., and J.M. Lund. 2015. Genetic diversity in the collaborative cross model recapitulates human West Nile virus disease outcomes. MBio 6:e00493–00415.

Gralinski, L.E., M.T. Ferris, D.L. Aylor, A.C. Whitmore, R. Green, M.B. Frieman, D. Deming, V.D. Menachery, D.R. Miller, R.J. Buus, T.A. Bell, G.A. Churchill, D.W. Threadgill, M.G. Katze, L. McMillan, W. Valdar, M.T. Heise, F. Pardo-Manuel de Villena, and R.S. Baric. 2015. Genome Wide Identification of SARS-CoV Susceptibility Loci Using the Collaborative Cross. PLoS Genet 11:e1005504.

Gralinski, L.E., V.D. Menachery, A.P. Morgan, A.L. Totura, A. Beall, J. Kocher, J. Plante, D.C. Harrison-Shostak, A. Schafer, F. Pardo-Manuel de Villena, M.T. Ferris, and R.S. Baric. 2017. Allelic Variation in the Toll-Like Receptor Adaptor Protein Ticam2 Contributes to SARS-Coronavirus Pathogenesis in Mice. G3 (Bethesda) 7:1653–1663.

Gralinski, L.E., T.P. Sheahan, T.E. Morrison, V.D. Menachery, K. Jensen, S.R. Leist, A. Whitmore, M.T. Heise, and R.S. Baric. 2018. Complement Activation Contributes to Severe Acute Respiratory Syndrome Coronavirus Pathogenesis. MBio 9:

Grifoni, A., D. Weiskopf, S.I. Ramirez, J. Mateus, J.M. Dan, C.R. Moderbacher, S.A. Rawlings, A. Sutherland, L. Premkumar, R.S. Jadi, D. Marrama, A.M. de Silva, A. Frazier, A.F. Carlin, J.A. Greenbaum, B. Peters, F. Krammer, D.M. Smith, S. Crotty, and A. Sette. 2020. Targets of T Cell Responses to SARS-CoV-2 Coronavirus in Humans with COVID-19 Disease and Unexposed Individuals. Cell 181:1489–1501 e1415.

Jameson, S.C. 2005. T cell homeostasis: keeping useful T cells alive and live T cells useful. Seminars in immunology 17:231–237.

Keane, T.M., L. Goodstadt, P. Danecek, M.A. White, K. Wong, B. Yalcin, A. Heger, A. Agam, G. Slater, M. Goodson, N.A. Furlotte, E. Eskin, C. Nellaker, H. Whitley, J. Cleak, D. Janowitz, P. Hernandez-Pliego, A. Edwards, T.G. Belgard, P.L. Oliver, R.E. McIntyre, A. Bhomra, J. Nicod, X. Gan, W. Yuan, L. van der Weyden, C.A. Steward, S. Bala, J. Stalker, R. Mott, R. Durbin, I.J. Jackson, A. Czechanski, J.A. Guerra-Assuncao, L.R. Donahue, L.G. Reinholdt, B.A. Payseur, C.P. Ponting, E. Birney, J. Flint, and D.J. Adams. 2011. Mouse genomic variation and its effect on phenotypes and gene regulation. Nature 477:289–294.

Kim, J.M., J.P. Rasmussen, and A.Y. Rudensky. 2007. Regulatory T cells prevent catastrophic autoimmunity throughout the lifespan of mice. Nat Immunol 8:191–197.

Kollmus, H., C. Pilzner, S.R. Leist, M. Heise, R. Geffers, and K. Schughart. 2018. Of mice and men: the host response to influenza virus infection. Mamm Genome 29:446–470.

Lanteri, M.C., K.M. O’Brien, W.E. Purtha, M.J. Cameron, J.M. Lund, R.E. Owen, J.W. Heitman, B. Custer, D.F. Hirschkorn, L.H. Tobler, N. Kiely, H.E. Prince, L.C. Ndhlovu, D.F. Nixon, H.T. Kamel, D.J. Kelvin, M.P. Busch, A.Y. Rudensky, M.S. Diamond, and P.J. Norris. 2009. Tregs control the development of symptomatic West Nile virus infection in humans and mice. J Clin Invest 119:3266–3277.

Lanzer, K.G., T. Cookenham, W.W. Reiley, and M.A. Blackman. 2018. Virtual memory cells make a major contribution to the response of aged influenza-naive mice to influenza virus infection. Immun Ageing 15:17.

Le Campion, A., C. Bourgeois, F. Lambolez, B. Martin, S. Leaument, N. Dautigny, C. Tanchot, C. Penit, and B. Lucas. 2002. Naive T cells proliferate strongly in neonatal mice in response to self-peptide/self-MHC complexes. Proceedings of the National Academy of Sciences of the United States of America 99:4538–4543.

Lee, D.C., J.A. Harker, J.S. Tregoning, S.F. Atabani, C. Johansson, J. Schwarze, and P.J. Openshaw. 2010. CD25+ natural regulatory T cells are critical in limiting innate and adaptive immunity and resolving disease following respiratory syncytial virus infection. J Virol 84:8790–8798.

Lee, J.Y., S.E. Hamilton, A.D. Akue, K.A. Hogquist, and S.C. Jameson. 2013. Virtual memory CD8 T cells display unique functional properties. Proceedings of the National Academy of Sciences of the United States of America 110:13498–13503.

Leist, S.R., and R.S. Baric. 2018. Giving the Genes a Shuffle: Using Natural Variation to Understand Host Genetic Contributions to Viral Infections. Trends Genet 34:777–789.

Loebbermann, J., H. Thornton, L. Durant, T. Sparwasser, K.E. Webster, J. Sprent, F.J. Culley, C. Johansson, and P.J. Openshaw. 2012. Regulatory T cells expressing granzyme B play a critical role in controlling lung inflammation during acute viral infection. Mucosal Immunol 5:161–172.

Lucas, C., P. Wong, J. Klein, T.B.R. Castro, J. Silva, M. Sundaram, M.K. Ellingson, T. Mao, J.E. Oh, B. Israelow, T. Takahashi, M. Tokuyama, P. Lu, A. Venkataraman, A. Park, S. Mohanty, H. Wang, A.L. Wyllie, C.B.F. Vogels, R. Earnest, S. Lapidus, I.M. Ott, A.J. Moore, M.C. Muenker, J.B. Fournier, M. Campbell, C.D. Odio, A. Casanovas-Massana, I.T. Yale, R. Herbst, A.C. Shaw, R. Medzhitov, W.L. Schulz, N.D. Grubaugh, C. Dela Cruz, S. Farhadian, A.I. Ko, S.B. Omer, and A. Iwasaki. 2020. Longitudinal analyses reveal immunological misfiring in severe COVID-19. Nature

Lund, J.M., L. Hsing, T.T. Pham, and A.Y. Rudensky. 2008. Coordination of early protective immunity to viral infection by regulatory T cells. Science 320:1220–1224.

Mateus, J., A. Grifoni, A. Tarke, J. Sidney, S.I. Ramirez, J.M. Dan, Z.C. Burger, S.A. Rawlings, D. M. Smith, E. Phillips, S. Mallal, M. Lammers, P. Rubiro, L. Quiambao, A. Sutherland, E. D. Yu, R. da Silva Antunes, J. Greenbaum, A. Frazier, A.J. Markmann, L. Premkumar, A. de Silva, B. Peters, S. Crotty, A. Sette, and D. Weiskopf. 2020. Selective and cross-reactive SARS-CoV-2 T cell epitopes in unexposed humans. Science

Mathew, D., J.R. Giles, A.E. Baxter, D.A. Oldridge, A.R. Greenplate, J.E. Wu, C. Alanio, L. Kuri-Cervantes, M.B. Pampena, K. D’Andrea, S. Manne, Z. Chen, Y.J. Huang, J.P. Reilly, A.R. Weisman, C.A.G. Ittner, O. Kuthuru, J. Dougherty, K. Nzingha, N. Han, J. Kim, A. Pattekar, E.C. Goodwin, E.M. Anderson, M.E. Weirick, S. Gouma, C.P. Arevalo, M.J. Bolton, F. Chen, S.F. Lacey, H. Ramage, S. Cherry, S.E. Hensley, S.A. Apostolidis, A.C. Huang, L.A. Vella, U.P.C.P. Unit, M.R. Betts, N.J. Meyer, and E.J. Wherry. 2020. Deep immune profiling of COVID-19 patients reveals distinct immunotypes with therapeutic implications. Science

McDermott, J.E., H.D. Mitchell, L.E. Gralinski, A.J. Eisfeld, L. Josset, A. Bankhead, 3rd, G. Neumann, S.C. Tilton, A. Schafer, C. Li, S. Fan, S. McWeeney, R.S. Baric, M.G. Katze, and K.M. Waters. 2016. The effect of inhibition of PP1 and TNFalpha signaling on pathogenesis of SARS coronavirus. BMC Syst Biol 10:93.

Min, B., R. McHugh, G.D. Sempowski, C. Mackall, G. Foucras, and W.E. Paul. 2003. Neonates support lymphopenia-induced proliferation. Immunity 18:131–140.

Pattacini, L., J.M. Baeten, K.K. Thomas, T.R. Fluharty, P.M. Murnane, D. Donnell, E. Bukusi, A. Ronald, N. Mugo, J.R. Lingappa, C. Celum, M.J. McElrath, J.M. Lund, and E.P.S.T. Partners Pr. 2016. Regulatory T-Cell Activity But Not Conventional HIV-Specific T-Cell Responses Are Associated With Protection From HIV-1 Infection. J Acquir Immune Defic Syndr 72:119–128.

Pruijssers, A.J., A.S. George, A. Schafer, S.R. Leist, L.E. Gralinksi, K.H. Dinnon, 3rd, B.L. Yount, M.L. Agostini, L.J. Stevens, J.D. Chappell, X. Lu, T.M. Hughes, K. Gully, D.R. Martinez, A.J. Brown, R.L. Graham, J.K. Perry, V. Du Pont, J. Pitts, B. Ma, D. Babusis, E. Murakami, J.Y. Feng, J.P. Bilello, D.P. Porter, T. Cihlar, R.S. Baric, M.R. Denison, and T.P. Sheahan. 2020. Remdesivir Inhibits SARS-CoV-2 in Human Lung Cells and Chimeric SARS-CoV Expressing the SARS-CoV-2 RNA Polymerase in Mice. Cell Rep 32:107940.

Qin, C., L. Zhou, Z. Hu, S. Zhang, S. Yang, Y. Tao, C. Xie, K. Ma, K. Shang, W. Wang, and D.S. Tian. 2020. Dysregulation of immune response in patients with COVID-19 in Wuhan, China. Clin Infect Dis

Rasmussen, A.L., A. Okumura, M.T. Ferris, R. Green, F. Feldmann, S.M. Kelly, D.P. Scott, D. Safronetz, E. Haddock, R. LaCasse, M.J. Thomas, P. Sova, V.S. Carter, J.M. Weiss, D.R. Miller, G.D. Shaw, M.J. Korth, M.T. Heise, R.S. Baric, F.P. de Villena, H. Feldmann, and M.G. Katze. 2014. Host genetic diversity enables Ebola hemorrhagic fever pathogenesis and resistance. Science 346:987–991.

Richert-Spuhler, L.E., and J.M. Lund. 2015. The Immune Fulcrum: Regulatory T Cells Tip the Balance Between Pro- and Anti-inflammatory Outcomes upon Infection. Prog Mol Biol Transl Sci 136:217–243.

Roberts, A., D. Deming, C.D. Paddock, A. Cheng, B. Yount, L. Vogel, B.D. Herman, T. Sheahan, M. Heise, G.L. Genrich, S.R. Zaki, R. Baric, and K. Subbarao. 2007a. A mouse-adapted SARS-coronavirus causes disease and mortality in BALB/c mice. PLoS Pathog 3:e5.

Roberts, A., F. Pardo-Manuel de Villena, W. Wang, L. McMillan, and D.W. Threadgill. 2007b. The polymorphism architecture of mouse genetic resources elucidated using genome-wide resequencing data: implications for QTL discovery and systems genetics. Mamm Genome 18:473–481.

Ruckwardt, T.J., K.L. Bonaparte, M.C. Nason, and B.S. Graham. 2009. Regulatory T cells promote early influx of CD8+ T cells in the lungs of respiratory syncytial virus-infected mice and diminish immunodominance disparities. J Virol 83:3019–3028.

Sariol, A., and S. Perlman. 2020. Lessons for COVID-19 Immunity from Other Coronavirus Infections. Immunity

Schuler, T., G.J. Hammerling, and B. Arnold. 2004. Cutting edge: IL-7-dependent homeostatic proliferation of CD8+ T cells in neonatal mice allows the generation of long-lived natural memory T cells. Journal of immunology 172:15–19.

Sheahan, T.P., A.C. Sims, R.L. Graham, V.D. Menachery, L.E. Gralinski, J.B. Case, S.R. Leist, K. Pyrc, J.Y. Feng, I. Trantcheva, R. Bannister, Y. Park, D. Babusis, M.O. Clarke, R.L. Mackman, J.E. Spahn, C.A. Palmiotti, D. Siegel, A.S. Ray, T. Cihlar, R. Jordan, M.R. Denison, and R.S. Baric. 2017. Broad-spectrum antiviral GS-5734 inhibits both epidemic and zoonotic coronaviruses. Sci Transl Med 9:

Smigiel, K.S., S. Srivastava, J.M. Stolley, and D.J. Campbell. 2014. Regulatory T-cell homeostasis: steady-state maintenance and modulation during inflammation. Immunol Rev 259:40–59.

Soerens, A.G., A. Da Costa, and J.M. Lund. 2016. Regulatory T cells are essential to promote proper CD4 T-cell priming upon mucosal infection. Mucosal Immunol 9:1395–1406.

Sosinowski, T., J.T. White, E.W. Cross, C. Haluszczak, P. Marrack, L. Gapin, and R.M. Kedl. 2013. CD8alpha+ dendritic cell trans presentation of IL-15 to naive CD8+ T cells produces antigen-inexperienced T cells in the periphery with memory phenotype and function. Journal of immunology 190:1936–1947.

Surh, C.D., and J. Sprent. 2005. Regulation of mature T cell homeostasis. Seminars in immunology 17:183–191.

Weiskopf, D., K.S. Schmitz, M.P. Raadsen, A. Grifoni, N.M.A. Okba, H. Endeman, J.P.C. van den Akker, R. Molenkamp, M.P.G. Koopmans, E.C.M. van Gorp, B.L. Haagmans, R.L. de Swart, A. Sette, and R.D. de Vries. 2020. Phenotype and kinetics of SARS-CoV-2-specific T cells in COVID-19 patients with acute respiratory distress syndrome. Sci Immunol 5:

Welsh, C.E., D.R. Miller, K.F. Manly, J. Wang, L. McMillan, G. Morahan, R. Mott, F.A. Iraqi, D.W. Threadgill, and F.P. de Villena. 2012. Status and access to the Collaborative Cross population. Mamm Genome 23:706–712.

Wilk, A.J., A. Rustagi, N.Q. Zhao, J. Roque, G.J. Martinez-Colon, J.L. McKechnie, G.T. Ivison, T. Ranganath, R. Vergara, T. Hollis, L.J. Simpson, P. Grant, A. Subramanian, A.J. Rogers, and C.A. Blish. 2020. A single-cell atlas of the peripheral immune response in patients with severe COVID-19. Nat Med 26:1070–1076.

Zhao, J., A.N. Alshukairi, S.A. Baharoon, W.A. Ahmed, A.A. Bokhari, A.M. Nehdi, L.A. Layqah, M.G. Alghamdi, M.M. Al Gethamy, A.M. Dada, I. Khalid, M. Boujelal, S.M. Al Johani, L. Vogel, K. Subbarao, A. Mangalam, C. Wu, P. Ten Eyck, S. Perlman, and J. Zhao. 2017. Recovery from the Middle East respiratory syndrome is associated with antibody and T-cell responses. Sci Immunol 2:

Zhao, J., J. Zhao, A.K. Mangalam, R. Channappanavar, C. Fett, D.K. Meyerholz, S. Agnihothram, R.S. Baric, C.S. David, and S. Perlman. 2016. Airway Memory CD4(+) T Cells Mediate Protective Immunity against Emerging Respiratory Coronaviruses. Immunity 44:1379–1391.

